# Single cell multiomic analysis of T cell exhaustion *in vitro*

**DOI:** 10.1101/846048

**Authors:** Mirko Corselli, Suraj Saksena, Margaret Nakamoto, Woodrow E. Lomas, Ian Taylor, Pratip K. Chattopadhyay

## Abstract

A key step in the clinical production of CAR-T cells is the expansion of engineered T cells. To generate enough cells for a therapeutic product, cells must be chronically stimulated, which raises the risk of inducing T-cell exhaustion and reducing therapeutic efficacy. As protocols for T-cell expansion are being developed to optimize CAR T cell yield, function and persistence, fundamental questions about the impact of *in vitro* manipulation on T-cell identity are important to answer. Namely: 1) what types of cells are generated during chronic stimulation? 2) how many unique cell states can be defined during chronic stimulation? We sought to answer these fundamental questions by performing single-cell multiomic analysis to simultaneously measure expression of 39 proteins and 399 genes in human T cells expanded *in vitro*. This approach allowed us to study – with unprecedented depth - how T cells change over the course of chronic stimulation. Comprehensive immunophenotypic and transcriptomic analysis at day 0 enabled a refined characterization of T-cell maturational states (from naïve to TEMRA cells) and the identification of a donor-specific subset of terminally differentiated T-cells that would have been otherwise overlooked using canonical cell classification schema. As expected, T-cell activation induced downregulation of naïve-associated markers and upregulation of effector molecules, proliferation regulators, co-inhibitory and co-stimulatory receptors. Our deep kinetic analysis further revealed clusters of proteins and genes identifying unique states of activation defined by markers temporarily expressed upon 3 days of stimulation (PD-1, CD69, *LTA)*, markers constitutively expressed throughout chronic activation (CD25, GITR, *LGALS1*), and markers uniquely up-regulated upon 14 days of stimulation (CD39, *ENTPD1, TNFDF10*). Notably, different ratios of cells expressing activation or exhaustion markers were measured at each time point. These data indicate high heterogeneity and plasticity of chronically stimulated T cells in response to different kinetics of activation. In this study, we demonstrate the power of a single-cell multiomic approach to comprehensively characterize T cells and to precisely monitor changes in differentiation, activation and exhaustion signatures in response to different activation protocols.

## Introduction

The heterogeneity of T-cells is remarkable; many genes and proteins (“markers”) are associated with cell maturity, trafficking, activation, and function, and these markers can be dramatically modulated over the course of an immune response(Chattopadhyay and Chiu, 2019; Chattopadhyay and Roederer, 2012; Chattopadhyay et al., 2019). Many are dysregulated by the tumor microenvironment(Nettey et al., 2018), so understanding their expression patterns provides critical insight(s) into biological mechanisms of disease, as well as information about potential drug targets. Moreover, it is likely that expression patterns of these markers may be useful in predicting disease, treatment outcome, or therapy-related adverse events(Spencer et al., 2016).

Technologies to measure T-cell associated markers have evolved dramatically in the past decade(Chattopadhyay et al., 2018; Chattopadhyay et al., 2019). The use of platforms that assay cells in bulk (like microarrays) has fallen out of favor, because these approaches average expression across many cells, even though the cells vary individually in expression. Single cell RNA sequencing (sc-RNAseq) overcomes the limitations of bulk measurements, and is powerful because of the large number of transcripts that can be interrogated simultaneously(Papalexi and Satija, 2018). However, transcription is a “noisy” process that occurs in bursts at irregular intervals and varies even across isogenic/clonal cells(Raj and van Oudenaarden, 2008), therefore cell populations often cannot be clearly resolved based on transcriptional analysis alone.

Cell subset discrimination is greatly enhanced, however, when protein and transcript analysis are combined(Stoeckius et al., 2017). Molecular cytometry, a new class of single cell technologies, adapts next generation sequencing (NGS) to single cell analysis to simultaneously provide information about cellular transcripts and proteins(Chattopadhyay et al., 2019). Like flow cytometry, cells are stained with antibodies and unbound antibodies are washed away before analysis. However, unlike flow cytometry, cells are labeled with oligonucleotide-tagged antibodies (rather than fluorescent molecules), and loaded onto an instrument that captures single cells and lyses them. Beads capture cellular mRNA as well as oligonucleotides associated with cell-bound antibodies via poly A-oligo dT interactions. Single cell sequencing then reveals the number of antibodies bound and the target of each bound antibody (as represented by a unique oligonucleotide tag). In addition, the number and identity of targeted cellular transcripts (or whole transcriptome) are also measured. In this manuscript, we demonstrate the use of molecular cytometry to measure the expression of 38 proteins and 399 targeted T cell transcripts that are relevant to immune cell biology, including T cell exhaustion.

Molecular cytometry technologies, like CITE-Seq(Stoeckius et al., 2017) REAP-Seq,^8^ and BD^TM^ AbSeq (AbSeq, presented here), carry a number of advantages over other single cell technologies. First, molecular cytometry platforms measure many more parameters than fluorescence or mass cytometry(Chattopadhyay et al., 2019) at the single cell level, allowing exquisitely detailed and comprehensive analysis of immune responses. Second, fluorescence and mass cytometry both require subtraction of signals that overlap across channels (compensation)(Chevrier et al., 2018; Cossarizza et al., 2019; Maciorowski et al., 2017); and this requirement poses significant challenges in terms of experimental design and data analysis. The oligonucleotide tags used in molecular cytometry are unique, which eliminates the need for cross-channel correction. Third, molecular cytometry can simultaneously interrogate cellular mRNA and proteins more easily than flow or mass cytometry, allowing study of post-transcriptional regulation of protein expression. In sum, molecular cytometry technologies offer deeper profiling of immune responses.

Upon antigenic challenge, immune responses are maintained and amplified by activated T-cells, which express unique transcriptional and protein signatures(Dominguez et al., 2017). In the case of an acute infection, markers associated with T cell activation drive key cellular processes, including proliferation, recruitment, homing, cytokine secretion, and cytotoxicity, ultimately resulting in resolution of the immune insult. However, upon sustained antigenic stimulation, as in the case of chronic viral infection or cancer, some activated T cells progressively develop an exhausted phenotype, which is characterized by reduced or lost effector function (e.g., loss of cytokine secretion) and impaired proliferation(Wherry and Kurachi, 2015a). During the course of an immune response, the degree to which these exhausted T-cells are generated dictates key outcomes, including whether a pathogen is cleared or an organism is chronically infected(Blank et al., 2019), the potential for responsiveness of a tumor to checkpoint inhibition therapy(Hashimoto et al., 2018), and potentially, the outcome of adoptive immunotherapy using chimeric antigen-receptor (CAR)-T cells(Ghoneim et al., 2016).

CAR-T cell therapy, in particular, offers a unique setting to explore the utility of molecular cytometry-based immune-profiling. A key step in the production of CAR-T cell clinical products is the expansion of genetically engineered T-cells using, among other methodologies, bead-coated antibodies specific for CD3 and CD28 and by treating the cells with exogenous IL2(Wang and Riviere, 2016). To generate enough engineered cells for therapeutic potency, cells must be chronically stimulated, which raises the risk that exhausted T-cells could dominate the infusion product, and in turn, reduce therapeutic efficacy. Although characterization of therapeutic products has direct patient relevance, more fundamental questions are also important to answer, namely: 1) what degree of heterogeneity and plasticity do T cells exhibit during chronic stimulation? 2) how many unique cell states (based on transcriptional and protein expression profiles) define chronic stimulation? and 3) what markers discriminate activated from exhausted cells?

We sought to answer these questions by using molecular cytometry to study an *in vitro* model system that resembles methods typically used to produce CAR T-cells. Our study revealed with unprecedented depth how T cells change upon chronic stimulation at the genotypic and phenotypic level, and also highlighted the use of molecular cytometry for immune profiling.

## Results

### Chronic Stimulation of CD8^+^ T cells *in vitro* Recapitulates Phenotypic and Functional Features of Exhaustion

We have developed two *in vitro* models mimicking either chronic or transient T-cell stimulation. Chronic stimulation was achieved by continuously stimulating T cells with recombinant human IL-2 (rhIL-2) and *α*CD3/CD28 beads for 14 days. Transient stimulation was achieved by stimulating T cells with rh-IL2 and *α*CD3/CD28 beads for 3 days, followed by resting in the presence of rhIL-2 for additional 11 days. Cells were collected at different time points, as indicated in Figure 1A.

**Figure 1.**
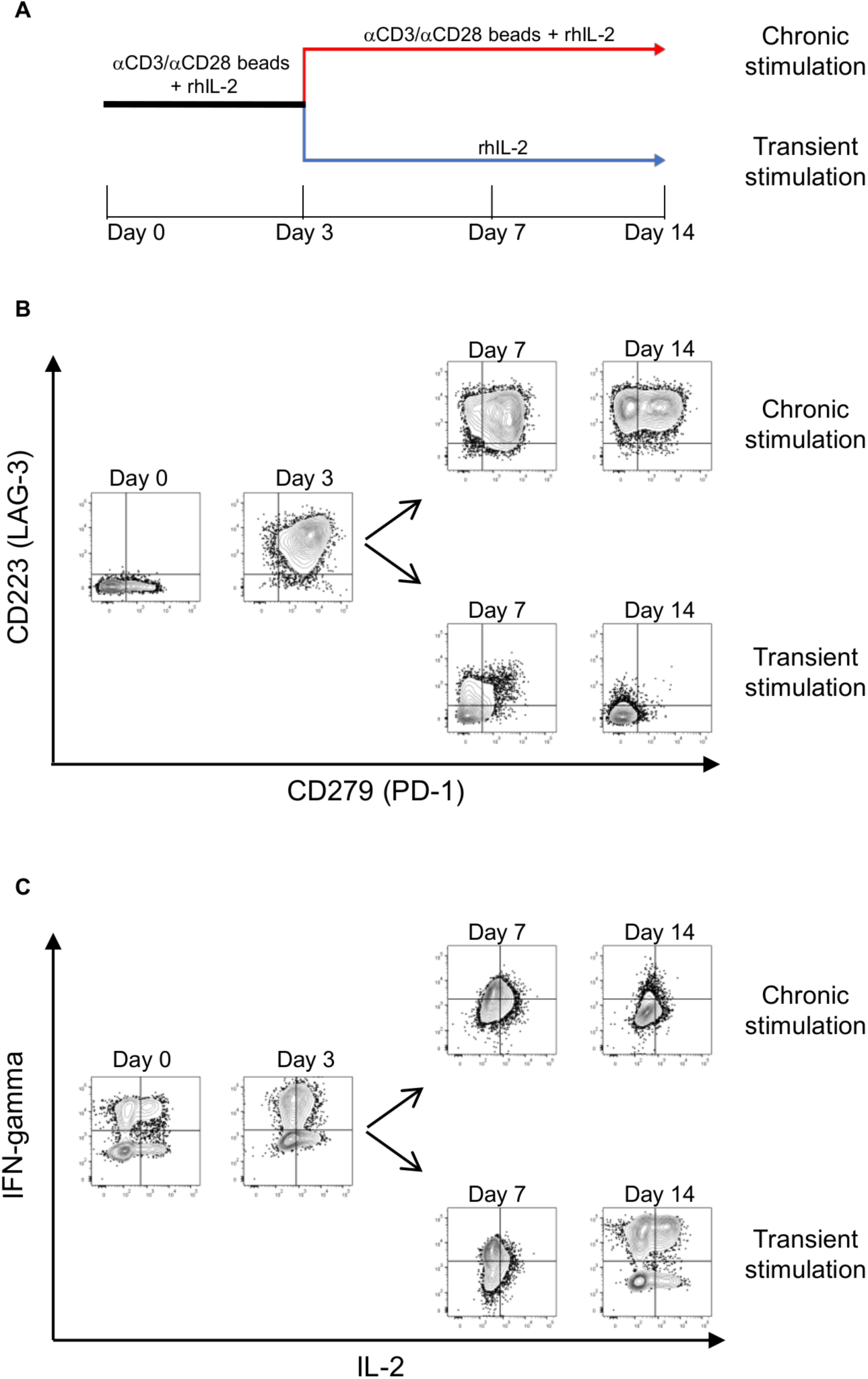
*In vitro* chronic stimulation recapitulates features of T-cell exhaustion. (A) Depiction of the *in vitro* model used for chronic and transient stimulation of T cells isolated from healthy donors. Cells were collected and frozen at Day 0, 3, 7 and 14 for downstream analysis. (B) Representative flow cytometry analysis of the expression of inhibitory receptors CD279 (PD-1) and CD223 (LAG-3) on CD8+ T cells in response to chronic or transient stimulation. (C) Representative intracellular flow cytometry analysis of CD8+ T-cell function in response to chronic or transient stimulation, followed by PMA/Ionomycin stimulation. Analyses performed on fresh cells from at least three independent experiments.

To assess whether CD8^+^ T cells in our *in vitro* model system acquired phenotypic and functional features characteristic of chronically stimulated T cells, we measured the upregulation of well-characterized inhibitory receptors, and the production of inflammatory cytokines using flow cytometry. CD8^+^ T cells demonstrated appreciable upregulation of the inhibitory receptors CD279 (PD-1) and CD223 (LAG-3) upon three days of stimulation (Figure 1B). LAG-3 expression was maintained through day 14 upon chronic stimulation, with a gradual downregulation of PD-1 for some cells, as previously described. Under transient stimulation conditions, CD8^+^ T cells progressively lost expression of both inhibitory receptors by day 14. Next, we analyzed T-cell function at each time point, and for each experimental condition (*i.e.,* chronic vs. transient stimulation). At baseline and after three days of culture with rhIL-2/*α*CD3/CD28, cells produced IFN*γ* and/or IL2; however, at day 7 – in both chronic and transient conditions – cytokine expression was greatly reduced (Figure 1C). This observation suggests that cells become functionally impaired beyond three days of stimulation. Cytokine production after 14 days was dramatically impaired when cells were stimulated continually (chronic stimulation), while T-cell function was recovered in transient stimulation conditions (Figure 1C). Together, these results demonstrate that well-established phenotypic and functional changes associated with T-cell exhaustion are specifically and gradually induced in our *in vitro* model system(Wherry and Kurachi, 2015b).

### AbSeq Enables Protein Detection with Specificity and Resolution Equivalent to Flow Cytometry

Having confirmed that our model system models the dynamics of marker expression associated with T cell exhaustion, we next compared the expression of different cell surface proteins, as measured by flow cytometry and AbSeq. The flow cytometry and AbSeq panels used for the comparison are outlined in Table 1. Figure 2A shows CD39 expression on total T cells (CD8+ and CD8-(mostly CD4+)) over the time course of the study. CD39 expression on CD8^+^ and CD8^-^ T-cells was detectable in a small subset of cells upon 7 days of chronic stimulation. After 14 days of stimulation, the frequency of CD8^+^ and CD8^-^ T cells expressing CD39 increased, along with the level of CD39 expression. In contrast, no CD39 expression was detected in any T-cell subset after 14 days of transient stimulation. Once detectable, CD39 expression patterns were qualitatively similar for both AbSeq- and flow cytometry-based measurements. For 7 of 10 proteins whose detection was compared using the two approaches, AbSeq and flow cytometry reported similar frequency of cells expressing each marker (Figure 2B & Supplemental Figure 1A). The lower frequency of LAG3^+^, CD62L^+^, and CTLA4^+^ observed by flow cytometry may be attributable to sub-optimal performance of the fluorochrome-conjugated antibodies, compared to the oligonucleotide conjugates. Additionally, the kinetics of marker expression were also very consistent between the flow cytometry and AbSeq measurements (Figure 2C & Supplemental Figure 1B). Thus, the sensitivity and specificity of the AbSeq approach are comparable to flow cytometry.

**Figure 2.**
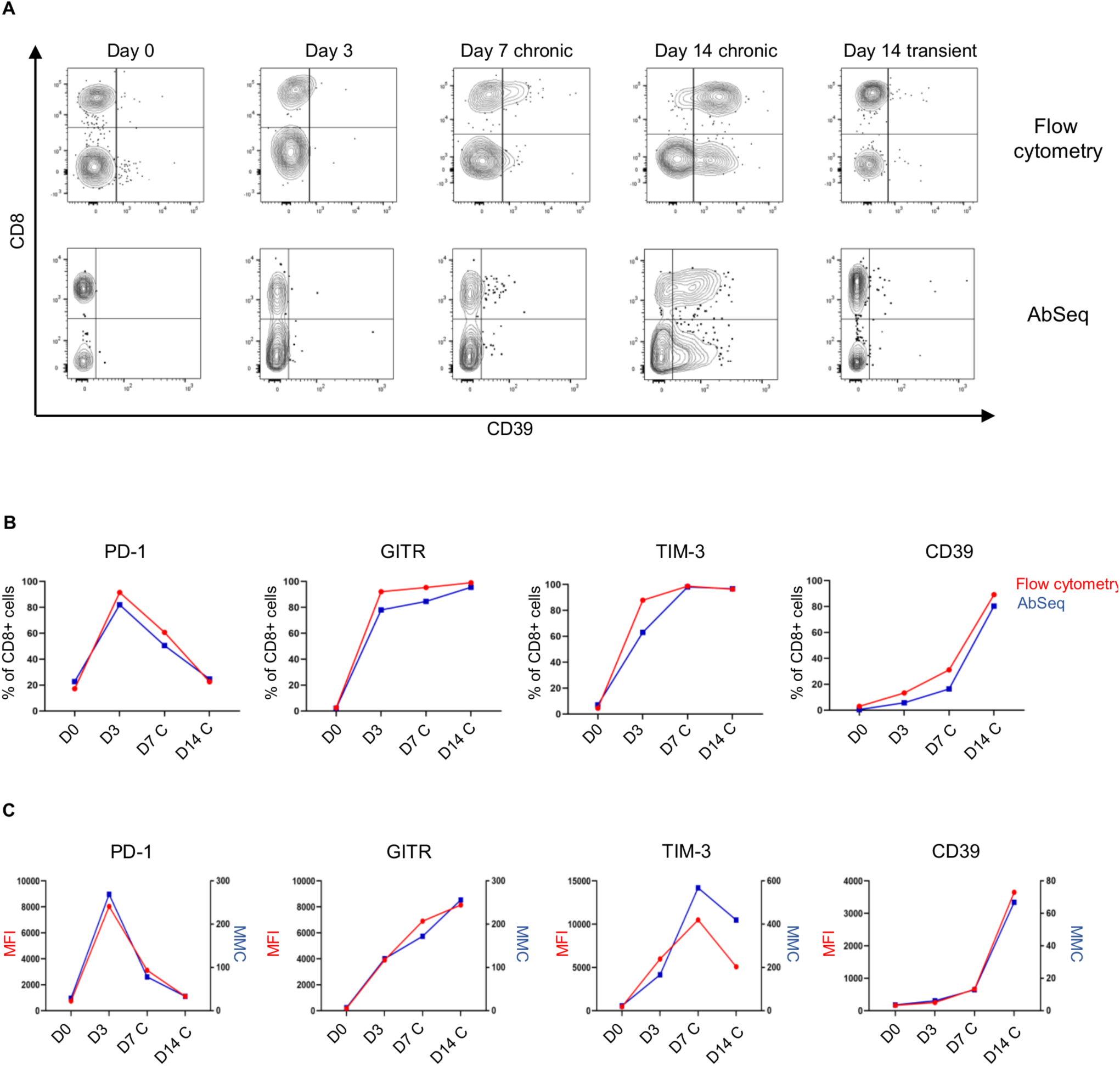
AbSeq and flow cytometry enable protein measurement with equivalent specificity and sensitivity. (A) Qualitative analysis of CD39 kinetic expression within CD8^+^ and CD8^-^ subsets of CD3^+^ T cells performed using either flow cytometry (top panel) or AbSeq (bottom panel). (B) Percentage of CD8^+^ cells expressing PD-1, GITR, TIM-3 and CD39 at day 0 (D0), day 3 (D3), day 7 (D7 C) and day 14 (D14 C) of chronic stimulation using either flow cytometry (red line) or AbSeq (blue line). (C) Levels of expression of PD-1, GITR, TIM-3 and CD39 on CD8^+^ T cells at day 0 (D0), day 3 (D3), day 7 (D7 C) and day 14 (Day14 C) of chronic stimulation. Mean fluorescence intensity (red line, left y axis) and mean molecular count (blue line, right y axis) were used to measure relative antigen expression levels using flow cytometry and AbSeq, respectively. This side-by-side analysis was performed on different aliquots of the same cells derived from the same donor (Donor 1), cryopreserved at the indicated time points. Flow cytometry data were down-sampled in order to analyze the same number of cells across the two platforms.

**Table 1.** Multiplex panels used for protein analysis. A 13-parameter panel was used for flow cytometry-based analysis. A 38-parameter panel was used for AbSeq-based analysis. Twelve specificities were common to both panels and were used to compare the performance of the two platforms.

| Marker | Clone | Flow Cytometry or AbSeq | Fluorochrome/Dye used for flow cytometry analysis |
| --- | --- | --- | --- |
| CD4 | SK3 | Both | BUV805 |
| CD8 | RPA-T8 | Both | BUV395 |
| CD45RA | HI100 | Both | APC-H7 |
| CD62L | DREG-56 | Both | FITC |
| CD95 | DX2 | Both | BV786 |
| CD279 (PD-1) | EH12.1 | Both | PE-Cy7 |
| CD223 (LAG-3) | T47-530 | Both | BV480 |
| CD366 (TIM-3) | 7D3 | Both | BV711 |
| CD357 (GITR) | V27-580 | Both | BV421 |
| CD152 (CTLA-4) | BNI3 | Both | PE |
| CD39 | TU66 | Both | BUV737 |
| CD103 | Ber-ACT8 | Both | APC |
| Live/Dead | N/A | Flow Cytometry Only | 7-AAD |
| CD3 | SK7 | AbSeq Only | N/A |
| CD14 | MφP-9 | AbSeq Only | N/A |
| B7-H4 | MIH43 | AbSeq Only | N/A |
| CD127 | HIL-7R-M21 | AbSeq Only | N/A |
| CD134 | ACT35 | AbSeq Only | N/A |
| CD137 | 4B4-1 | AbSeq Only | N/A |
| CD154 | TRAP1 | AbSeq Only | N/A |
| CD183 | 1C6/CXCR3 | AbSeq Only | N/A |
| CD185 | RF8B2 | AbSeq Only | N/A |
| CD194 | 1G1 | AbSeq Only | N/A |
| CD196 | 11A9 | AbSeq Only | N/A |
| CD197 | 3D12 | AbSeq Only | N/A |
| CD2 | RPA-2.10 | AbSeq Only | N/A |
| CD25 | 2A3 | AbSeq Only | N/A |
| CD27 | M-T271 | AbSeq Only | N/A |
| CD270 | CW10 | AbSeq Only | N/A |
| CD278 | DX29 | AbSeq Only | N/A |
| CD28 | CD28.2 | AbSeq Only | N/A |
| CD30 | BerH8 | AbSeq Only | N/A |
| CD38 | HIT2 | AbSeq Only | N/A |
| CD49a | SR84 | AbSeq Only | N/A |
| CD54 | HA58 | AbSeq Only | N/A |
| CD69 | FN50 | AbSeq Only | N/A |
| CD7 | M-T701 | AbSeq Only | N/A |
| CD94 | HP-3D9 | AbSeq Only | N/A |
| CD98 | UM7F8 | AbSeq Only | N/A |

### Molecular Cytometry Enables Deeper Profiling of Effector T Cells

Next, we conducted a multi-omic analysis of resting CD8+ T cells (day 0) by simultaneously measuring the expression of 38 proteins and 399 T cell-specific genes at the single cell level. The 38-plex AbSeq panel is outlined in Table 1; and the list of the 399 genes measured by the targeted RNAseq panel is reported in Supplemental file 1. First, we used AbSeq to quantify the different subsets of unstimulated CD8^+^ T cells (day 0) based on differential expression of markers commonly used for identification of T-cell differentiation states(Chattopadhyay and Roederer, 2010; Mahnke et al., 2013). CD45RA and CD28 were used to define CD45RA^-^ CD28^+^ central memory cells (CM), CD45RA^-^ CD28^-^ effector memory cells (EM) and CD45RA^+^ CD28^-^ effector memory RA cells (EMRA). CD27 was additionally used for the identification of CD45RA^+^ CD28^+^ CD27^high^ naïve cells (N). As expected, both qualitative and quantitative analysis revealed differences across the three donors when looking at the different T cell subsets (Figure 3A, B). We also noted a unique population of CD45RA^+^CD28^+^CD27^low^ cells in donor 3, which represented ∼12% of the CD8^+^ T cell compartment.

**Figure 3.**
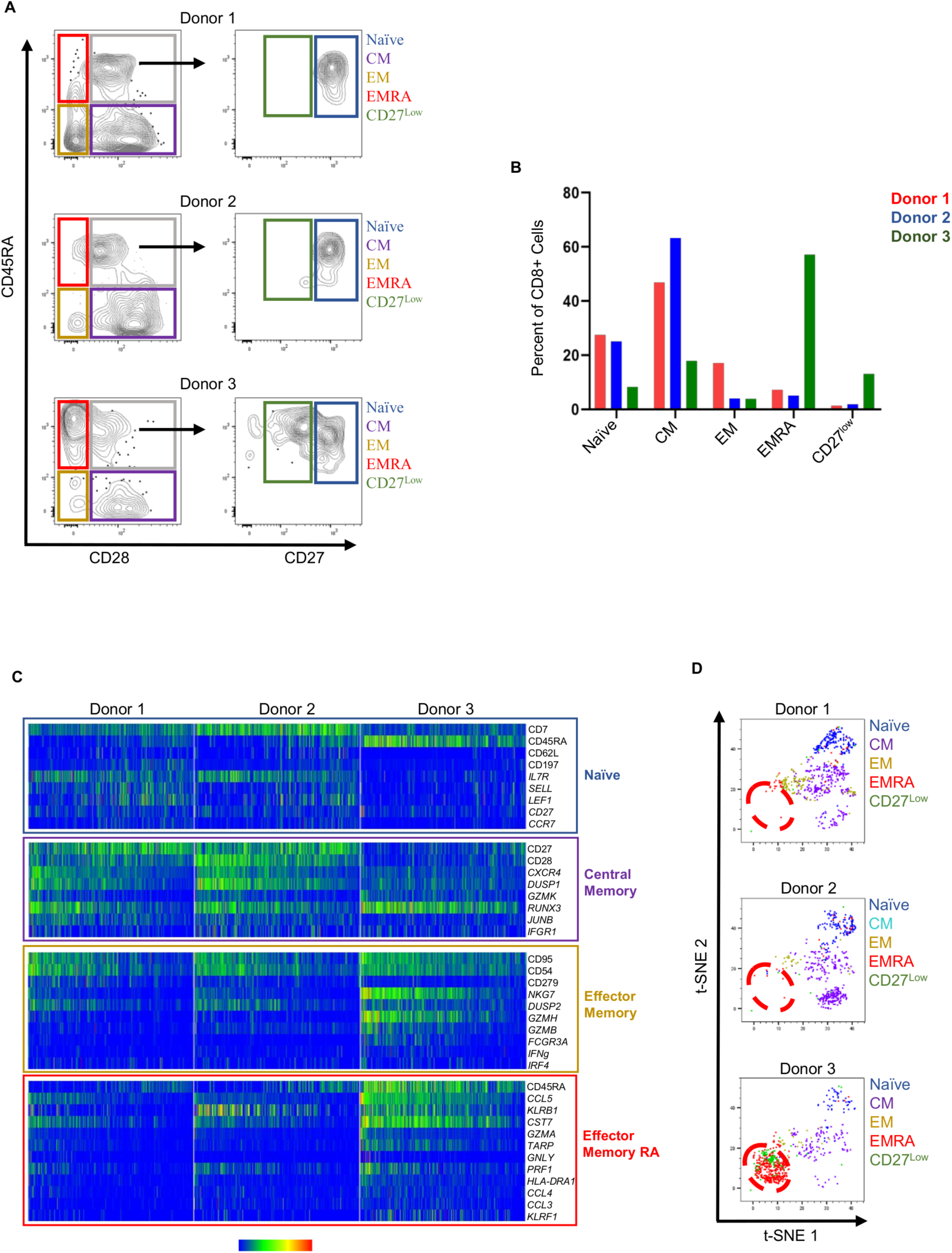
Deep characterization of fresh CD8+ T-cell maturational states. (A) Gating strategy used to identify CD8^+^ naïve (blue box), central memory (CM, purple box), effector memory (EM, yellow box), effector memory RA (EMRA, red box) T cells from 3 donors at day 0 based on measurement of CD45RA, CD28 and CD27 expression via AbSeq. A unique population of CD45RA^+^CD28^+^CD27^low^ cells (CD27^low^, green box) was detected in donor 3. (B) Frequency of CD8+ T-cell subsets across the three donors. (C) Single-cell heatmap of selected proteins and genes (*italics*) differentially expressed in each T-cell subset across the three donors (fold change ≥2, q≤0.05). Four hundred cells per donor are represented. Each column represents a single cell. Event columns are colored on a 0-100% min-max pseudocolor scale based on relative parameter expression. (D) t-SNE visualization of the CD8^+^ T-cell subsets showing different cell clusters with a naïve, CM, EM, EMRA and CD27^low/-^ phenotype across the three donors. T-SNE plots were generated based on expression of 38 proteins and highly dispersed genes. Cells were color-coded based on the phenotype described in panel A. The red dashed circle indicates EMRA and CD27^low/-^ cells clustering together in donor 3.

To further define proteins and transcripts associated with the different maturation states across donors, we performed differential gene and protein expression profiling comparing naïve cells to CM, EM, and EMRA T cells. Consistent with published findings(Mahnke et al., 2013), we observed higher levels of CD45RA, CD62L, CD197 (CCR7) proteins and *SELL, LEF1, IL7R and CCR7* transcripts in naïve cells (Supplemental file 2). Also, as expected, the multi-omic analysis revealed that EM and EMRA cells (yellow and red boxes) express high levels of PD-1 protein and transcripts encoding cytotoxic proteins NKG7 (*NKG7),* granzymes *(GZMA, GZMB, GZMH),* granulysin (*GNLY*), and perforin (*PRF1*) (Supplemental file). Interestingly, CD8^+^ T cells from donor 3 showed higher expression of effector-associated markers, compared to samples from donors 1 and 2 that were enriched for cells displaying a naïve/central memory profile (Figure 3C). Also, our analysis revealed new markers not previously associated with the different T cell maturation states (PIK3IP1, PASK, and TXK in CD8^+^ N cells and DUSP1 and IFNGR1 in CD8^+^ CM cells).

To better characterize the CD45RA^+^CD28^+^CD27^low^ population in Donor 3, we generated t-distributed stochastic neighbor embedding (t-SNE) maps for each donor. The events in the t-SNE plot are color-coded based on the maturation state of cells as defined by differential expression of CD45RA, CD28 and CD27 (Figure 3D). The high number of EMRA cells in Donor 3 formed a distinct cell cluster that included the CD27^low^ cells (Figure 3C, bottom panel, red dashed circle), suggesting that CD27^low^ cells share gene and protein expression patterns with EMRA cells rather than CD45RA^+^CD28^+^CD27^high^ naïve T-cells, as might have been expected. To test whether CD27^low^ cells are more closely associated with EMRA than naïve cells, we performed differential expression analysis, comparing all measured transcripts and proteins between CD27^low^ and naïve cells derived from Donor 3. We found that genes and their corresponding proteins common to naïve cells (and absent from EMRA), like *SELL/*CD62L, *IL7R/*CD127, *CCR7/*CD197, were expressed at higher levels in naïve cells compared to CD27^low^ cells (Supplemental Table 1). Conversely, CD27^low^ cells were enriched for proteins and genes expressed by EMRA cells, like PD-1, *PRF1*, *GZMB*, and *IFNG* (Supplemental Table 2). Notably, 19 of the 20 markers detected at higher levels in CD27^low^ cells (compared to naïve cells) were also enriched in EMRA cells (Supplemental Table 3). In sum, our results demonstrate that CD27^low^ cells, that might have been wrongly classified as naïve cells based solely on expression of CD45RA, CD28, and CD27, are in fact, likely to be a subset of effector memory cells capable of re-expression of both CD45RA and CD27. This analysis reveals the power of multi-omic (protein and mRNA) analyses for more precise identification of cell types, and for deeper profiling of classical T-cell phenotypes.

### Molecular Cytometry Identifies Gene and Protein Signatures Associated with Different Modes of T Cell Activation

Having confirmed the utility of molecular cytometry for detailed cellular profiling, next, we used this approach to investigate temporal changes in protein and gene expression that accompany chronic T-cell stimulation and exhaustion. The same AbSeq and targeted gene panels (Table 1 and Supplemental File 1) that were used for the characterization of T-cell maturational states, were used for a comprehensive analysis of CD8^+^ T cells at different stages of chronic and transient stimulation. High-dimensional datasets were generated for each donor at the different time points based on expression of 38 proteins and highly dispersed genes. Projection of cells from the of the three concatenated donors using t-SNE revealed three major cell groups (Figure 4A, “All Donors”). The first group encompassed events from resting cells (before activation, “Day 0,” red colored events), while the second group included events from 3, 7, and 14 days of chronic stimulation (blue, orange, and green colored events), and the third group comprised cells from the 3-day stimulation + 11-day rest condition (“Day 14 Transient,” purple colored events). These results demonstrate that expression changes among the 38 markers measured by AbSeq in conjunction with targeted gene expression analysis clearly capture the distinction between resting and activated cells.

**Figure 4.**
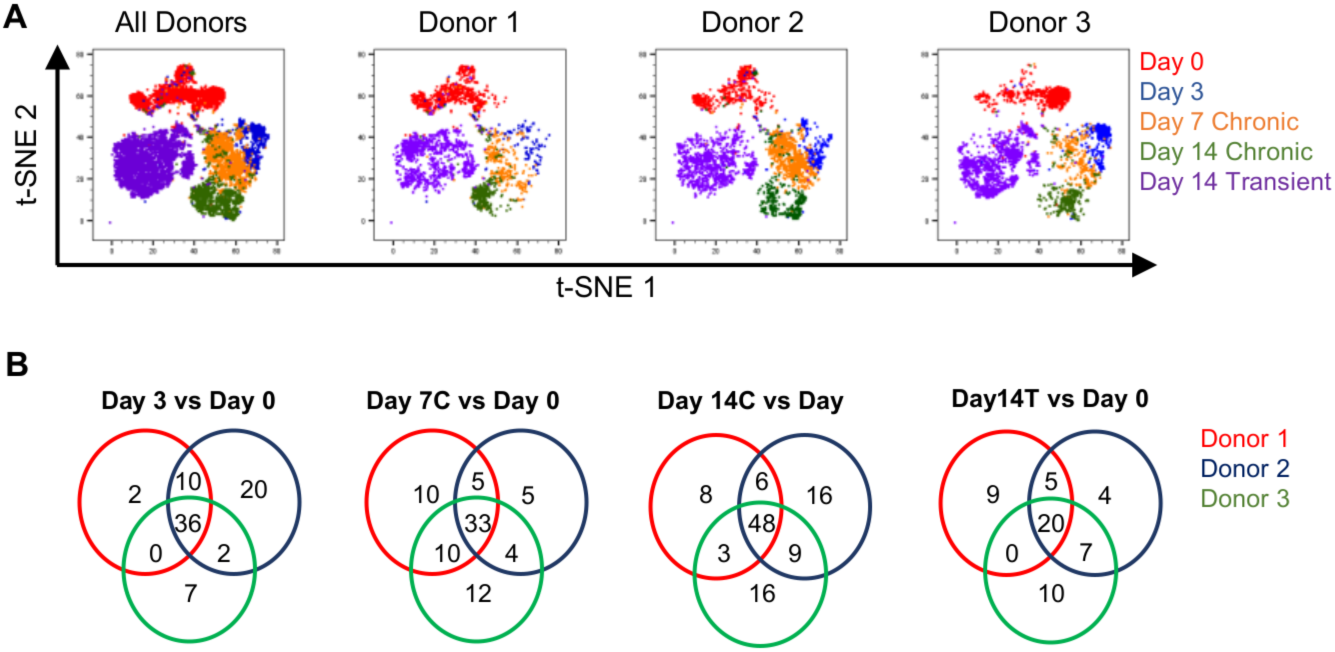
Identification of common signatures associated with T-cell activation. (A) t-SNE visualization of CD8+ cells clusters at day 0 (red), day 3 (blue), day 7 and 14 of chronic stimulation (orange and green, respectively), and day 14 of transient stimulation (purple). t-SNE plots were generated based on expression of 38 protein and highly dispersed genes. (B) Venn diagrams indicate the number of shared or uniquely upregulated genes and proteins across the three donors at day 3, day 7 chronic stimulation (Day 7C), day 14 chronic stimulation (D14C), and day 14 transient (D14T) stimulation, as compared to unstimulated cells (Day 0).

To identify the genes and proteins uniquely associated with the mode of T-cell activation (chronic versus transient), we performed differential expression analysis for each individual donor by comparing each time point of the *in vitro* activation system to each other (Supplemental File 3). By comparing the individual lists of differentially expressed genes and proteins, we identified, for example, common signatures defined by markers upregulated in each donor at each time point, as compared to unstimulated cells (Day 0). The results of this analysis are summarized using the Venn diagrams in Figure 4B. We also observed upregulation of genes and/or proteins shared by 2 out of 3 donors, or unique to each donor. The complete list of shared and unique markers upregulated at each time point (in each donor), as compared to Day 0, is reported in Supplemental File 4.

Next, we focused on genes and proteins that were upregulated 3-fold or higher upon activation in at least two of the three donors. This approach resulted in the identification of four sets of genes and proteins with unique expression patterns. The first set (Figure 5A, red bar) consisted of markers elevated after three days of stimulation (D3), but downregulated thereafter (CD278, CD69, *IFNg*, *IL9*, and *Lymphotoxin A* (*LTA*)). The second set (Figure 5A, blue) represented markers elevated after 14 days of chronic stimulation (D14C), and included *TNFSF10*, *YBX3*, *CSF2*, *BIRC3*, *ENTPD1*, and CD39. The third set of markers (Figure 5A, green) was upregulated at all stimulation time points, but generally downregulated when the stimulus was removed (as measured after three days stimulation and 11 days rest; D14T in Figure 5A). This set included genes and proteins that could be broadly classified into markers of activation/proliferation (CD25, CD357, CD54, CD98, CD137, *GZMB*, *IL2Ra*, *PCNA*, *TOP2A*, and *TYMS*) versus inhibition/exhaustion (CD223, *IRF4*, *LAG3*, *LGALS1*, and *ZBED2*). The fourth set (Figure 5A, purple) encompassed markers whose expression was upregulated in cells transiently stimulated. Taken together, our analysis reveals unique activation signatures that can be differentially linked to the duration of activation.

**Figure 5.**
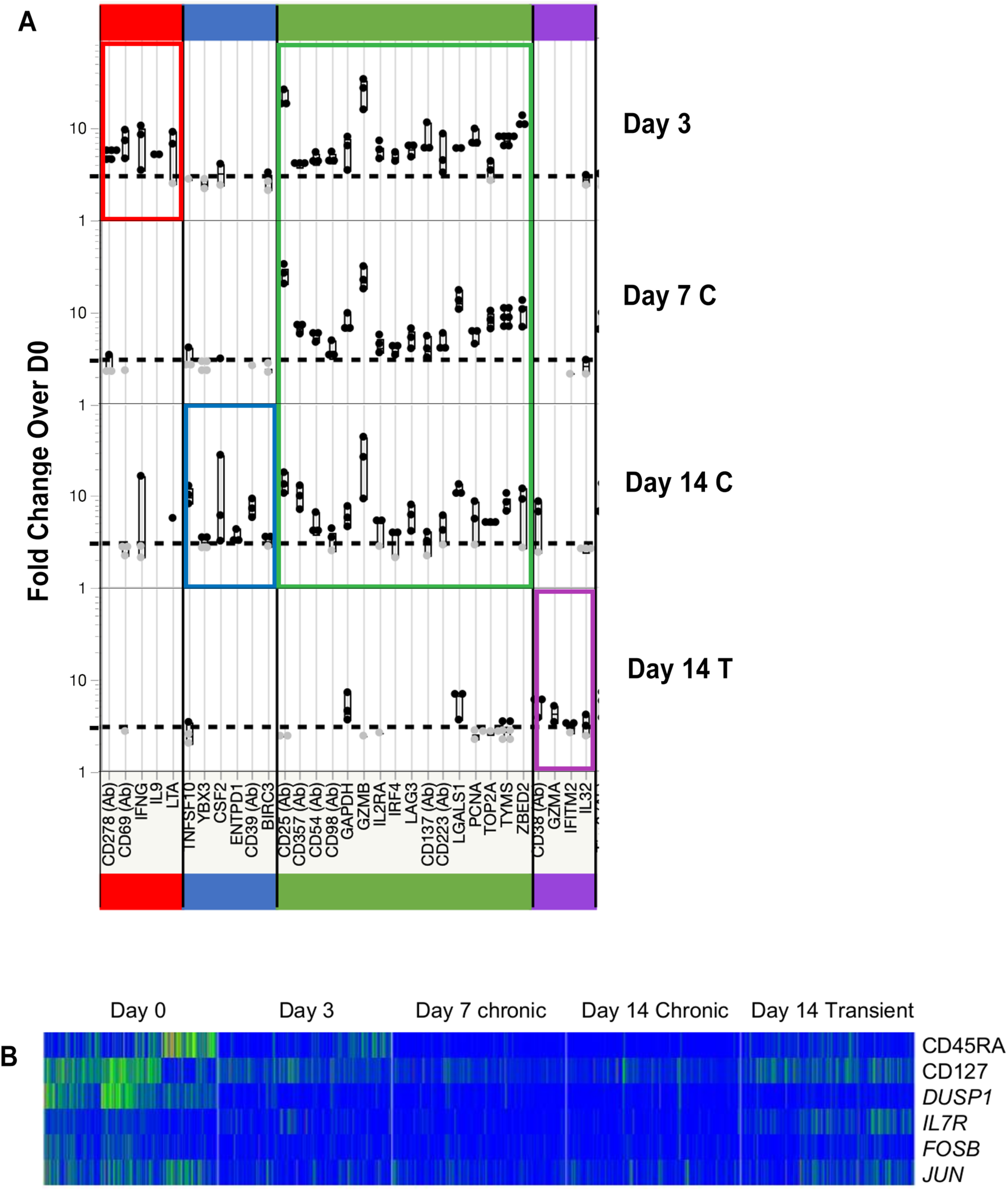
Identification signatures associated with distinct modes of T-cell activation. (A) Markers whose expression is upregulated (≥ three-fold, in at least two donors; left y-axis), as compared to day 0, at each time point (right y-axis). Markers uniquely upregulated with three days of stimulation are indicated by the red bar, while markers unique to chronic 14-day stimulation are indicated by the blue bar. The green bar indicates which markers are elevated at all time points in the chronically stimulated cells, while the purple bar denotes markers uniquely associated with the three day stimulation / 11 day rest (transient stimulation, D14T) condition. (B) Single-cell heatmap of proteins and genes (italic) upregulated in all three donors at Day 0 and downregulated upon cell activation (fold change ≥2, q≤0.05). Data from four hundred and twenty cells measured at each time point. An equal number of cells (140) from each of the three donors is represented for each time point.

Notably, our analysis also revealed a set of genes and proteins that were selectively associated with resting cells in all three donors. These markers were down-regulated upon stimulation, and then partially re-acquired after the stimulation was removed. This marker set included proteins and genes associated with naïve T-cells (CD45RA, *IL7R,* and its corresponding protein CD127), along with *DUSP1, FOSB* and *JUN* (Figure 5B).

### Molecular cytometry reveals unique relationships between inhibitory and proliferation markers

To understand whether markers of activation or inhibition are expressed on distinct cells or if these genes and proteins could be co-expressed on the same cells, we plotted expression in bivariate plots using concatenated data from the 3 donors. Amongst the various combinations of markers, those involving *LGALS1* and *ZBED2* transcripts stood out for their progressive expression over the time course. At baseline, very few cells expressed *LGALS1* (the mRNA for Galectin-1, an immune inhibitory molecule) or proliferation markers *TYMS* and *PCNA* (Figure 6A). At day 3, *LGALS1* and both proliferation markers were upregulated, and cells expressing all possible combinations of markers (i.e., one marker alone, neither marker, both markers) could be detected (Figure 6A). However, after 7 days of stimulation, almost all cells expressing *TYMS* or *PCNA* co-expressed the inhibitory molecule *LGALS1*, suggesting the inhibitory potential of these cells. Conversely, after only three days of stimulation, the great majority of cells expressing *TYMS* and *PCNA* co-expressed *ZBED2,* a transcription factor associated with progression to T-cell dysfunction (Figure 6B). Beyond three days, the frequency of cells expressing *ZBED2* was gradually reduced, with a concurrent increase in the frequency of cells single positive for *PCNA* and *TYMS* (Figure 6B). Thus, expression of LGALS1 and ZBED2 appear reciprocal over the time course, and the ratio of these two molecules may mark how long cells have been stimulated (Figure 6C).

**Figure 6.**
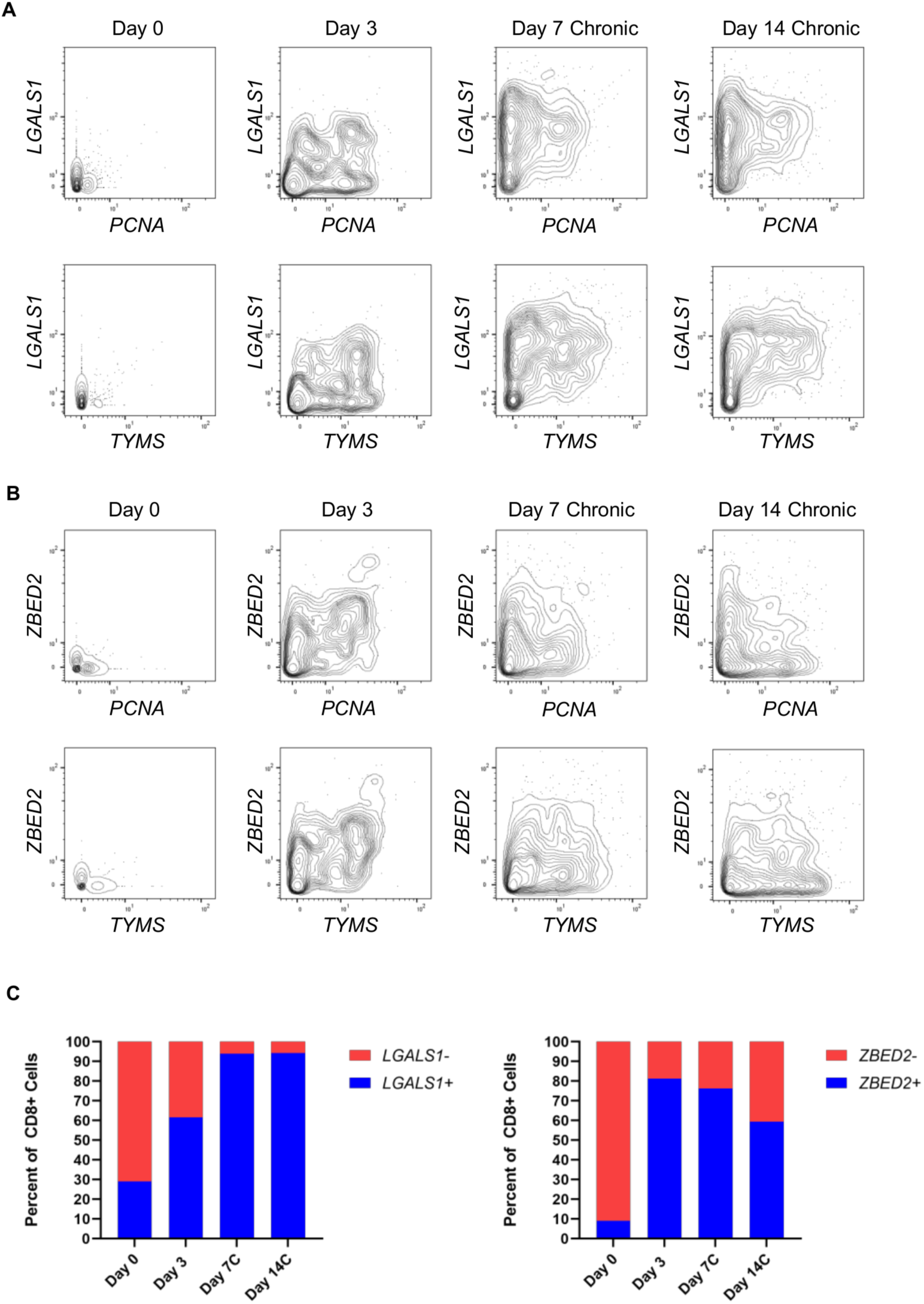
Combinatorial analysis of inhibitory and proliferation marker expression. Bi-variate plots showing the relationship between the inhibitory markers *LGALS1* (A) or *ZBED2* (B) and the proliferation markers *TYMS* or *PCNA* throughout chronic stimulation. (C) Frequency of CD8+ T-cells expressing *LGALS1* and *ZBED2* throughout chronic stimulation. Data were generated from concatenated samples from 3 donors.

### Molecular Cytometry Reveals Correlation Between Cellular Transcription & Translation

In addition to detailed cellular profiling, molecular cytometry technologies provide a unique ability to correlate the expression of genes and their corresponding proteins at the single-cell level. In this study, we examined this correlation for 23 pairs of genes and their corresponding proteins, and determined if kinetic changes (over the activation time course) were similar. In some cases, mRNA and the corresponding protein exhibited concordance in terms kinetics of expression. For example, *ENTPD1* and its corresponding protein CD39 were concordantly upregulated at day 14 of chronic stimulation (Figure 7A). Similarly, both *IL7R* and its corresponding protein CD127 were concordantly downregulated upon chronic cell stimulation. For certain markers, we observed discordant patterns for mRNA and protein expression. For example, unstimulated cells at day 0 express basal levels of *CD69* mRNA but not protein. Upon 3 days of stimulation, we observed *CD69* mRNA downregulation, with concomitant upregulation of CD69 protein. After 3 days of stimulation, both *CD69* mRNA and protein were downregulated. Finally, in some cases, protein expression was detected (and varied over the course of the stimulation), without concomitant measurable mRNA expression (*PDCD1* mRNA/PD1 protein, Figure 7A; other cases in Supplemental Figure 2).

**Figure 7.**
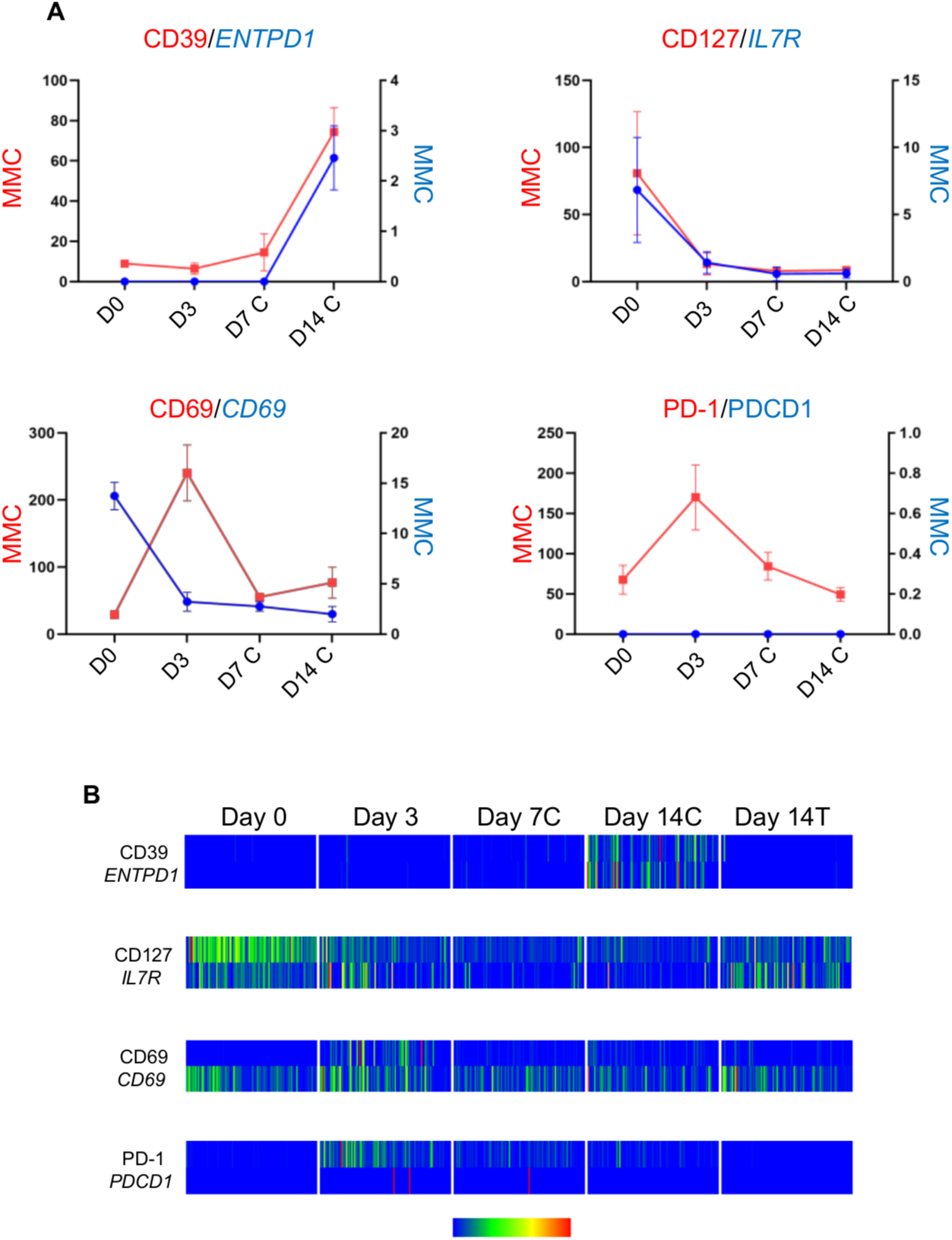
Correlation between gene and protein expression at the single cell level. (A) Kinetic analysis of mRNA and protein expression. mRNA and protein levels were measured as mean molecular count (MMC) and depicted by the red trace (protein) and blue trace (mRNA) at day 0 (D0), day 3 (D3), day 7 (D7 C) and day 14 (D14 C) of chronic stimulation. Analysis performed on three individual donors. Data represented as mean ± standard deviation. (B) Single-cell heatmap of gene and protein expression. A single cell is represented in each column. One hundred cells from a single donor (donor 2) are represented at each time point. Event columns are colored on a 0-100% min-max pseudocolor scale based on relative parameter expression.

Amongst the cases where no mRNA was detected, this result was largely driven by the inability to detect any transcript at any time point (*PDCD1*, *CD28*, *TNFRSF8*, *TNFRSF4*, and *TNFRSF9*), suggesting lack of expression or expression below limits of detection. In contrast, for CD69, CD27, and CD2, mRNA for a marker was present at detectable levels, but the kinetics of expression (the change between time points) was not similar for mRNA and protein. In sum, the relationship between protein and mRNA expression was complex, and observed in all possible patterns (concordant/discordant in frequency of expression; concordant/discordant in level of expression). This is consistent with previous reports describing the stochastic nature of mRNA transcription(Raj et al., 2006).

The above analyses summarize the data on protein/gene expression correlation, but do not illustrate the heterogeneity of mRNA/protein concordance on a cell-by-cell basis. Concordance and discordance at the single cell is shown by heat map in Figure 7B. These heat maps reveal the complexity of mRNA/protein expression and kinetics at the single-cell level. In accordance with the similar kinetics observed for *ENTPD1* and CD39 (both upregulated after 14 days of chronic stimulation; Figure 7A), the great majority of cells (represented by colored bars) co-expressed mRNA and protein (Figure 7B). The same was not true for *IL7R* and CD127. Despite the overall similarity in kinetics between *IL7R* and CD127 (expressed at day 0, reduced at day 3, almost off on days 7 and 14; Figure 7A), a higher degree of heterogeneity was observed, with variable numbers of cells expressing protein only, gene only or both over time. In contrast, *CD69* mRNA is expressed by many cells at all time points (Figure 7B), but protein is expressed at essentially a single time point (day 3). For the *PDCD1*/PD1 pair, we observe high levels of PDCD1 in at least three cells analyzed (Figure 7B), suggesting that lack of PDCD1 transcript might not reflect low assay sensitivity. These results further demonstrate the value of single cell analysis.

## Discussion

Molecular cytometry is a potentially powerful method for immune monitoring; however, the technology is relatively new. We sought to provide data that qualifies AbSeq, a new molecular cytometry technology, for use in immune monitoring and to demonstrate the power of the approach. We first confirmed the validity of AbSeq as a sensitive methodology for protein measurement, as compared to the gold standard flow cytometry. As past studies(Peterson et al., 2017; Stoeckius et al., 2017) have shown, the proportion of cells expressing various markers are similar across the two platforms. In fact, when protein expression is visualized using bivariate plots, and as expected when using the same antibody clone for the two different methods, the patterns are quite similar across the technologies. Where we found discordance in the percentage of cells expressing markers such as LAG-3, CD62L, and CTLA-4, we suspect that this was a function of poor performance of the fluorochrome-conjugated antibody, since it is well-recognized that performance of an antibody clone can depend on the fluorochrome chosen or the design of a multicolor panel(Maciorowski et al., 2017). This issue highlights two important advantages of molecular cytometry over flow cytometry. First, all antibody tags are oligonucleotide sequences, so they share the same chemical and biophysical properties. As such, antibody performance in molecular cytometry is not dependent on the tag chosen or labeling method. Second, signal, background, and sensitivity are similar for all antibody tags that are sequenced, unlike flow cytometry, where the detectors (i.e., photomultiplier tubes) used for detection of each fluorochrome can vary a great deal(Perfetto et al., 2014). Moreover, in flow cytometry, the signals observed for a given level of protein expression may differ by fluorochrome, whereas the digital nature of molecular cytometry signals allow a consistent relationship to protein expression. In principle, quantification of receptor levels by molecular cytometry will therefore be considerably more straightforward than flow cytometry, requiring few (if any) standards to translate oligonucleotide molecules to relative receptor expression.

Unlike other molecular cytometry publications, we measured kinetics of marker expression, which provide further evidence of the concordance between flow and molecular cytometry. Our kinetic analysis also allowed comparison between protein and transcript expression for the 23 mRNAs with a coordinate antibody target. We found great variety in the relationships between transcript and protein expression amongst individual markers; we did not find high correlation between mRNA and protein expression, and often observed transcripts that were up-regulated or present without concomitant protein expression, and vice versa. In some cases, discordance may reflect protein or transcript expression that is below the level of detection. However, we often found – for phenotypically similar cells – that one cell had high levels of transcript while another cell did not. The relationship between transcript and protein expression did not follow any clear rules; it was not specific to proteins or cell types that shared a function, nor were there common kinetic patterns. This suggests that discordance between transcript and protein expression may reflect a) the stochastic nature of transcript expression, which occurs in bursts(Harper et al., 2011; Suter et al., 2011; Zhdanov, 2007) or b) regulation of transcript expression through mechanisms that are specific to individual proteins. In either case, it is very clear that transcript expression is not a direct surrogate for protein expression, nor is the abundance of an mRNA correlated with protein(Liu et al., 2016); these factors could diminish the value of platforms that solely measure single cell RNA expression in isolation (without consideration of protein expression) for immune monitoring.

Our analysis of naïve, memory, and effector T-cell subsets demonstrates the power of molecular cytometry for accurate identification of cell populations. In our oldest study subject (Donor 3, 61 years old), we observed a population of CD45RA^+^ cells with intermediate expression of CD27. Using canonical cell classification schema(Mahnke et al., 2013), such cells might have been categorized as naïve; however, by more deeply profiling cells – measuring the many proteins and genes available in our experiment – it became clear that CD27^low^ cells were much more similar to effector T-cells than naïve T-cells. The broader lesson revealed by this example is that molecular cytometry, and other high-parameter single-cell technologies can check and challenge the classical schema used to categorize the maturational status of cells. The problem presented by unusual patients or cell populations, which may be outliers in immune monitoring studies, can also be mitigated by molecular cytometry. The highly multiplexed, and simultaneous measurement of protein and mRNA, provides increased context that better describes unusual cell populations and allows more accurate classification and enumeration.

Our study examined, with great depth, how T cells change with activation, using molecular cytometry to analyze time course specimens from an *in vitro* activation model. The model recapitulated the process of T-cell activation and exhaustion(Wherry and Kurachi, 2015b), as confirmed by flow cytometric analysis of checkpoint molecules and cytokines. We identified sets of proteins and genes that are uniquely upregulated (compared to resting cells) at each time point we analyzed. Five markers were upregulated only at the three-day time point, including the genes coding for known activation marker CD69 and the effector cytokine interferon gamma. We posit that these markers, whose expression during short-term, *ex vivo* stimulation (6-24 hours) is well-documented by flow cytometry(Pitsios et al., 2008), remain elevated even after three days. Notably, expression of *IL9* and *lymphotoxin A* (LTA) mRNA were also highly upregulated at this time point. These represent new targets for immune assessment, perhaps using intracellular cytokine staining (ICS) by flow cytometry (provided that these mRNA are translated into protein). The identification of new immune monitoring targets – beyond the common cytokines measured by ICS – demonstrates the value of molecular cytometry.

We also identified the proteins and genes upregulated throughout the time course, which fell neatly into two groups – those that are associated with enhancing cell function and those that inhibit cell activity. Unlike bulk assays, in which thousands of cells are averaged for analysis, single cell molecular cytometry data allowed us to ask whether the upregulation of these markers was associated with two distinct cell types (activated vs. exhausted, for example) or whether these markers could be co-expressed. We found that many activation and inhibition markers were co-expressed, suggesting great plasticity in cell state and function, and that the relationships between some of these markers changed over the stimulation period. For example, markers associated with cell proliferation *PCNA* and *TYMS* were expressed by cells after three days of stimulation, with and without *LGALS1* expression. However, by day 7 of our stimulation assays, all cells expressing proliferation antigens co-expressed *LGALS1*. This gene transcript is notable because it encodes the Galectin-1 protein, which is known to inhibit cell proliferation in the tumor microenvironment and is a target of immunotherapy agents(Chou et al., 2018). Our result suggests the possible existence of an autocrine or paracrine feedback loop regulating cell proliferation, involving *PCNA* /*TYMS* and Galectin-1. Disruption of this feedback loop, using antibodies to Galectin-1, may provide a new means to prevent exhaustion of CAR-T cells during their manufacture. Similarly, the ability of CAR-T cells to persist *in vivo* might be enhanced by genetically engineering the over-expression of markers that are normally downregulated with stimulation, including *DUSP1, FOSB* and *JUN* (which were part of our resting cell signature). Indeed, a recent report describing “exhaustion-resistant” CAR-T cells with over-expressed c-Jun supports this possibility(Lynn et al., 2019). Finally, measurements of *JUN*, *LGALS1*, or the suite of markers upregulated with 14 days of stimulation may provide a predictor or indicator of the capacity of a CAR-T cell product to persist *in vivo*, raising the intriguing possibility of a companion diagnostic for CAR-T cell therapy.

The high parameter data provided by single cell molecular cytometry offers an unparalleled tool to better define a molecule’s function, expression, and disease relevance. For example, our study also reports reciprocal expression of *LGALS1* and *ZBED2*, in relationship to the expression of proliferation markers *PCNA* and *TYMS*; expression of *LGALS1* is gained in proliferating cells over stimulation, while *ZBED2* is lost from proliferating cells over the course of stimulation. *ZBED2* expression has recently been shown to mark a subset of melanoma-infiltrating CD8^+^ T cells poised to progress to a dysfunctional, exhausted state(Li et al., 2019). The same study also described highest and lowest proliferative potential at early and late stages of exhaustion, respectively, thus corroborating the importance of simultaneously assessing inhibitory and proliferation markers at the single-cell level.

We have also identified a set of markers exclusively expressed upon 14 days of chronic stimulation. Among these markers are the gene *ENTPD1* and its corresponding protein CD39, recently described as a marker defining tumor antigen-specific, exhausted TIL (Canale et al., 2018). The significant loss of cytokine production observed by flow cytometry with chronic stimulation suggests that expression of this set of markers correlates with T-cell dysfunction. Further studies investigating the expression of the identified signatures of primary tumor infiltrating lymphocytes are required to validate these hypotheses.

Ultimately, the power of molecular cytometry lies in the rich datasets it provides, which can be produced relatively easily – without the complications of experimental design found in other cytometry technologies. The technology approach offers an important advance in our ability to characterize – and exploit - cellular immunity.

## Materials and methods

### Cell preparation and cryopreservation

Peripheral blood was collected from N=4 healthy donors (age 28, 32, 36 and 61 years old) in accordance with approved IRB protocol BDX-ASCP0. Peripheral blood mononuclear cells (PBMCs) were isolated using Ficoll-Paque Plus (GE Healthcare). T cells were isolated using BD IMag^TM^ Human T Lymphocyte Enrichment Set, as per manufacturer’s instructions (BD Biosciences). Aliquots of cells collected at different time points of stimulation (day 0, 3, 7 and 14) were cryopreserved in freezing medium containing 90% fetal bovine serum (FBS; Hyclone Laboratories Inc.) and 10% DMSO (Sigma-Aldrich) and stored in liquid nitrogen. Cryopreserved cells were thawed in a 37°C water bath, diluted with 1ml of warm complete culture medium, then transferred to a tube containing 10ml of warm complete culture medium. Cells were centrifuged at 300g for 5 minutes prior to downstream processing. Cell viability of fresh or thawed cells was overall consistently ≥90%.

### T-cell stimulation

Freshly isolated T cells were plated in a 24-multiwell plate (Corning) at the concentration of 1×10^6^ cells/well in 2ml of complete culture medium composed of RPMI 1640 medium (Gibco) supplemented with 10% FBS, 1% Penicillin-Streptomycin (Hyclone Laboratories Inc.) and 1% L-glutamine (Hyclone Laboratories Inc.). To mimic a chronic stimulation, cells were cultured for 14 days in a humified CO_2_ incubator at 37°C in complete culture medium with Dynabeads® Human T-Activator CD3/CD28 beads (ThermoFisher Scientific) (25µl/well; bead-to-cell ratio of 1:1) and recombinant human interleukin-2 (rhIL-2; 25U/ml; Sigma-Aldrich). To mimic a transient stimulation, cells were cultured in the presence of CD3/CD28 beads and rhIL-2 for 3 days and then rested in the presence of rhIL-2 only for the remaining 11 days of culture. For both stimulation conditions, cells were collected at day 3, 7, 10 and 14. After collection, beads were magnetically removed using a BD IMag^TM^ Cell Separation Magnet (BD Biosciences) prior to cell preparation for flow cytometry analysis, passaging, cryopreservation. For cell passaging, cells were resuspended in fresh complete medium for chronic or transient stimulation and re-plated at the same cell density as at Day 0.

### Flow cytometry

Fresh or thawed T cells were resuspended in 2ml of BD Pharmingen^TM^ Stain Buffer (FBS; BD Biosciences) and then centrifuged at 300g for 5 minutes. For cell surface marker staining, 0.5-1×10^6^ cells were incubated for 30 minutes at 37°C with antibody cocktails composed of 100µl of Stain Buffer (FBS) (BD Biosciences), 10µl of BD Horizon^TM^ Brilliant Stain Buffer Plus (BSB; BD Biosciences) and each antibody at its recommended concentration, unless otherwise stated. Cells were incubated at room temperature (RT) for 30 minutes and then washed twice with Stain Buffer (FBS). Cells were resuspended in 0.5ml of Stain Buffer (FBS) and incubated with the viability dye 7-AAD (BD Biosciences) at RT for 10 minutes prior to acquisition on a 3-laser, 12-color BD FACSLyric^TM^ Research System or a 5-laser, 18-color BD LSRFortessa^TM^ X-20 Research Use Only system. For intracellular cytokine detection, the collected cells were first stimulated for 4 hours with phorbol 12-myristate 13-acetate (PMA; 50ng/ml; Sigma-Aldrich) and Ionomycin (500ng/ml; Sigma-Aldrich) in the presence of the transporter inhibitors BD GolgiPlug^TM^ and GolgiStop^TM^, as per manufacurer’s instructions (BD Biosciences). Cells were then washed with Stain Buffer (FBS) prior to surface marker staining, as per the protocol described above. Cells were then washed in Phosphate Buffered Saline (PBS) without FBS and stained with Fixable Viability Stain 620 (FVS620; BD Biosciences) as per manufacturer’s instructions. After 2 washes in Stain Buffer (FBS), cells were fixed and permeabilized using BD Cytofix/Cytoperm^TM^ Fixation/Permeabilization Solution, as per manufacturer’s instructions (BD Biosciences). Cells were then incubated at RT for 30 minutes with the antibody cocktail composed of 100µl of permeabilization buffer, 10µl of BSB and each antibody at recommended concentration, unless otherwise stated. Cells were then washed twice with permeabilization buffer, resuspended in 0.5ml of Stain Buffer (FBS) and acquired on a 3-laser, 12-color BD FACSLyric^TM^ Research System. After acquisition, all data were exported as FCS3.1 files and analyzed using FlowJo^TM^ software (version 10.6, BD Biosciences). The following mouse anti-human antibodies, all provided by BD Biosciences, were used in this study: CD4 APC-H7 (clone RPA-T4), CD4 APC-R700 (clone SK3), CD4 BUV805 (clone SK3), CD8 APC-H7 (clone SK1), CD8 AlexaFluor® 700 (clone RPA-T8), CD8 BUV395 (RPA-T8), CD279 PE-Cy7 (clone EH12.1), CD223 BV480 (LAG-3; clone T47-530) CD223 AlexaFluor® 647 (LAG-3; clone T47-530. 0.5µg/test), CD45RA APC-H7 (clone HI100), CD62L FITC (clone DREG-56), CD95 BV786 (clone DX2. 0.5µg/test), CD366 BV711 (TIM-3; clone 7D3), CD357 BV421 (GITR; clone V27-580), CD152 PE (CTLA-4; clone BNI3), CD39 BUV737 (clone Tu66), CD103 APC (clone Ber-Act8), interferon gamma FITC (IFN-gamma; clone B27), interleukin-2 PE (IL-2; clone MQ1-17H12).

### Single Cell Labelling with Sample Tags and AbSeq

Cell surface staining was performed as described in the protocol “Single Cell Labelling with the BD Single-Cell Multiplexing Kit and BD AbSeq Ab-oligos” (BD Biosciences). Briefly, cryopreserved T cells from 3 donors, were thawed as per the protocol described in cell processing section. To enable all samples (N=5 time points) for each donor to be loaded on a single BD Rhapsody^TM^ cartridge, the BD^TM^ Human Single-Cell Multiplexing kit (BD Biosciences) was used to label the cells from each donor with unique sample tag. Cells were sequentially labelled with sample tags followed by BD^TM^ AbSeq antibody-oligos (Ab-oligos). First, 1 million cells from each donor/stimulation condition were transferred to a vial containing a unique sample tag barcode (per donor) and incubated at room temperature for 20 minutes. Following incubation, cells were washed 3 times with Stain Buffer (FBS). Cells were counted and pooled together at an equal ratio of all conditions for each donor. A panel of Ab-oligos, described in Table 1, was prepared and added to the tube of 1 million pooled cells from each donor. Cells were incubated on ice for 30 minutes. Following incubation, cells were washed 3 times with Stain Buffer (FBS) and resuspended in Sample Buffer (FBS).

### Single Cell Capture and cDNA Synthesis

Cell capture was performed as described in the protocol “Single Cell Capture and cDNA Synthesis with the BD Rhapsody Single-Cell Analysis System” (BD Biosciences), using both the BD Rhapsody^TM^ Scanner and Express instrument. The BD Rhapsody scanner was used to perform cell count and viability using Calcein AM (Thermo Fisher Scientific) and Draq7 (BD. Biosciences). 20,000 pooled cells from each donor were loaded into 3 separate BD Rhapsody cartridges followed by cell capture beads. Cells were lysed and the capture beads were then retrieved and washed. Reverse transcription, followed by Exonuclease I treatment was performed on the retrieved cell capture beads, following manufacturer’s instructions.

### Library preparation

After undergoing cell capture and reverse transcription Rhapsody beads were taken into library preparation as described in “mRNA Targeted, Sample Tag, and BD AbSeq Library Preparation with the BD Rhapsody Targeted mRNA and AbSeq Amplification Kit”. Briefly, all beads were amplified in PCR1 using the BD Human Immune Response Panel (399 amplicons) + SMK, for 11 PCR cycles. Post-PCR1 reaction cleanup utilized a double-sided 0.7x/1.2x Ampure method to separate the larger mRNA PCR products from the smaller AbSeq/sample tag PCR products. Each of the Sample tag and mRNA products were taken into separate PCR2 reactions that utilize universal (SMK) or nested (mRNA) primers. 1.2x and 0.8x single-sided Ampure cleanups, respectively, were performed on the PCR2 products. AbSeq products were taken directly into indexing PCR after PCR1. mRNA PCR2 products (diluted to 1.4-2.7 ng/µL) and AbSeq PCR1/sample tag PCR2 products (diluted to 1.1 ng/µL) were taken into a 6-cycle indexing PCR reaction. SMK, mRNA, and AbSeq libraries from the same donor were indexed with the same reverse primer, with distinct indexes used between donors. Indexing PCR products for mRNA and AbSeq SMK utilized 0.7x and 0.8x single-sided Ampure cleanups, respectively. Indexing PCR reactions for mRNA and AbSeq/SMK All libraries were quantified using Agilent High Sensitivity DNA Analysis kits.

### Sequencing

All libraries were diluted to 2nM before pooling for sequencing. Preliminary sequencing for quality assessment was performed on an Illumina NextSeq 500 using a High Output 150 cycle kit with 75×75 bp PE reads. Libraries were pooled at a ratio of 1:5:12.5 (sample tag:mRNA:AbSeq) targeting 400 reads/cell from sample tag libraries, 2,000 reads/cell from mRNA libraries, and 5,000 reads/cell from AbSeq libraries. Full sequencing was done on an Illumina NovaSeq 6000 using an S1 kit with 75×75 bp PE reads. For the NovaSeq run the libraries were pooled at a ratio of 1:13:60 targeting an additional 150 reads/cell for sample tag libraries, 2,000 reads/cell for mRNA libraries, and 9,000 reads/cell from AbSeq libraries. Sequencing metrics are reported in Supplemental table 4.

### Bioinformatics analysis

FASTQ files were downloaded from Illumina BaseSpace and uploaded onto the Seven Bridges website. Each sample was run separately through the BD Rhapsody Analysis Pipeline using fastqs from the AbSeq panel and Human Immune Response Panel and using the “Single-Cell Multiplex Kit – Human” multiplexing setting. Output files in csv format were imported into SeqGeq v1.5 software (BD Biosciences) for AbSeq and scRNA-Seq data analysis.

### Dimensionality Reduction

In order to overcome visualization artifacts associated with sparse data, dimensionality reduction was performed using *principal component analysis* (PCA) guided *t-distributed stochastic neighbor embedding* (tSNE). The Opt-SNE optimized tSNE calculation was used to automatically detect and implement appropriate settings for this machine learning step in analysis(Belkina et al., 2019). Principal component analysis was performed on highly dispersed gene parameters in combination with extra-cellular antibody parameters detected via BD’s™ AbSeq pipeline.

### Differential Expression Analysis

Differential expression analysis was performed by pairwise comparisons in volcano plots; illustrating log2 fold change vs adjust p-Values, also known as “q-Values”, for differentially expressed genes (DEG). Mann-Whitney U-tests were utilized to estimate the reproducibility of observations in non-parametric distributions(Schurch et al., 2016). False Discovery Rate (FDR) adjusted p-Values were appropriate for clusters greater than 200 events in size. Inclusion criteria for DEG: fold-change values +/- 2.0 (up and down regulated, respectively) and q-values < 0.05.

### Heat-Maps

Single-cell heatmap figures were generated by first downsampling populations to a representative number of events. Event columns were then colored on a 0-100% min-max pseudocolor scale based on relative parameter expression, annotated in descending order.

## Tables

**Supplemental table 1.**
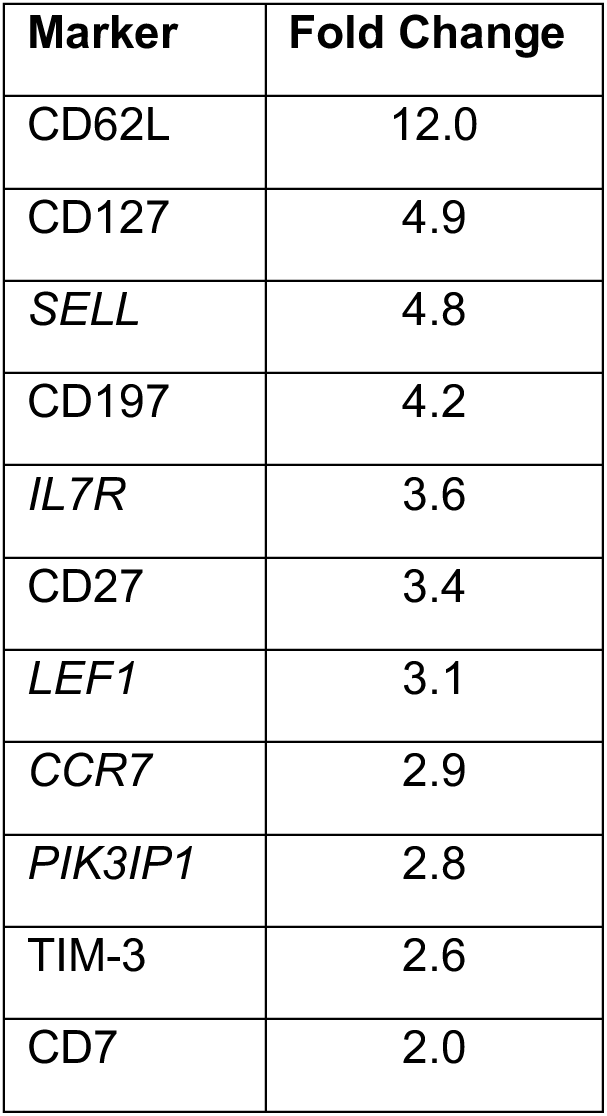
List of proteins and genes (italic) upregulated in CD8^+^CD45RA^+^CD28^+^CD27^high^ naïve cells, as compared to CD8^+^CD45RA^+^CD28^+^CD27^low^ cells. (fold-change ≥2, q value ≤0.05).

| Marker | Fold Change |
| --- | --- |
| CD62L | 12.0 |
| CD127 | 4.9 |
| *SELL* | 4.8 |
| CD197 | 4.2 |
| *IL7R* | 3.6 |
| CD27 | 3.4 |
| *LEF1* | 3.1 |
| *CCR7* | 2.9 |
| *PIK3IP1* | 2.8 |
| TIM-3 | 2.6 |
| CD7 | 2.0 |

**Supplemental table 2.** List of proteins and genes (italic) upregulated in CD8^+^CD45RA^+^CD28^+^CD27^low^ cells, as compared to CD8^+^CD45RA^+^CD28^+^CD27^high^ naïve cells. (fold-change ≥2, q value ≤0.05)

| Marker | Fold Change |
| --- | --- |
| *KLRB1* | 10.3 |
| *NKG7* | 8.2 |
| CD94 | 8.0 |
| *GZMH* | 7.1 |
| *CST7* | 7.0 |
| *CCL5* | 6.1 |
| *TARP* | 4.3 |
| *GZMA* | 4.1 |
| *FCGR3A* | 3.2 |
| *KLRF1* | 3.1 |
| *CD74* | 3.0 |
| *GZMK* | 3.0 |
| *DUSP2* | 2.7 |
| CD54 | 2.6 |
| *PRF1* | 2.5 |
| CD95 | 2.5 |
| *HLA-DPA1* | 2.3 |
| *CCL3* | 2.3 |
| *IL2RB* | 2.2 |
| *GZMB* | 2.2 |

**Supplemental table 3:**
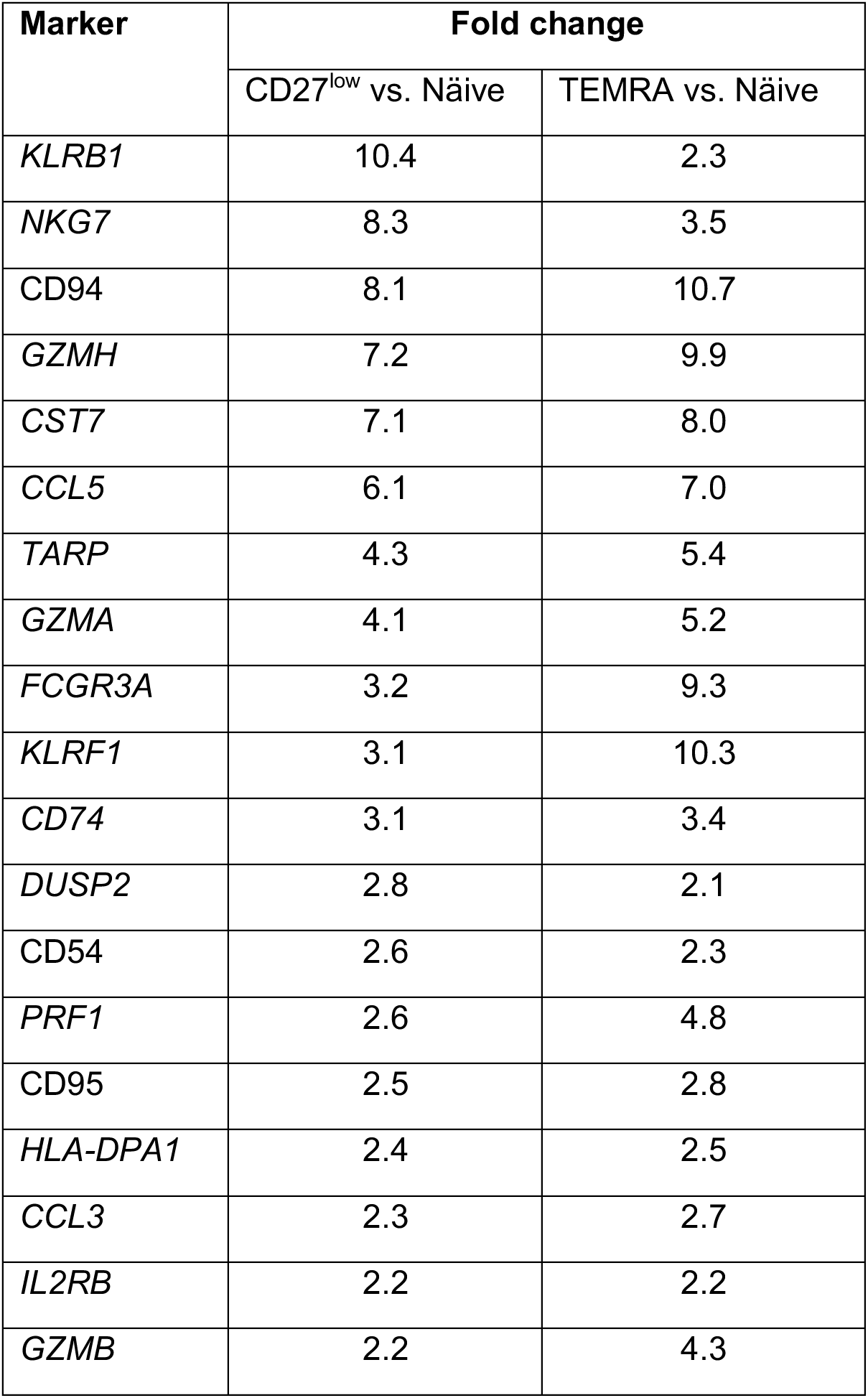
List of shared proteins and genes (italic) upregulated in CD8^+^CD45RA^+^CD28^+^CD27^low^ and CD8^+^CD45RA^+^CD28^-^ TEMRA cells, as compared to naïve cells. (fold change ≥2, q≤0.05).

| Marker | Fold change |  |
| --- | --- | --- |
|  | CD27<sup>low</sup> vs. Näive | TEMRA vs. Näive |
| *KLRB1* | 10.4 | 2.3 |
| *NKG7* | 8.3 | 3.5 |
| *CD94* | 8.1 | 10.7 |
| *GZMH* | 7.2 | 9.9 |
| *CST7* | 7.1 | 8.0 |
| *CCL5* | 6.1 | 7.0 |
| *TARP* | 4.3 | 5.4 |
| *GZMA* | 4.1 | 5.2 |
| *FCGR3A* | 3.2 | 9.3 |
| *KLRF1* | 3.1 | 10.3 |
| *CD74* | 3.1 | 3.4 |
| *DUSP2* | 2.8 | 2.1 |
| *CD54* | 2.6 | 2.3 |
| *PRF1* | 2.6 | 4.8 |
| *CD95* | 2.5 | 2.8 |
| *HLA-DPA1* | 2.4 | 2.5 |
| *CCL3* | 2.3 | 2.7 |
| *IL2RB* | 2.2 | 2.2 |
| *GZMB* | 2.2 | 4.3 |

**Supplemental Table 4.**
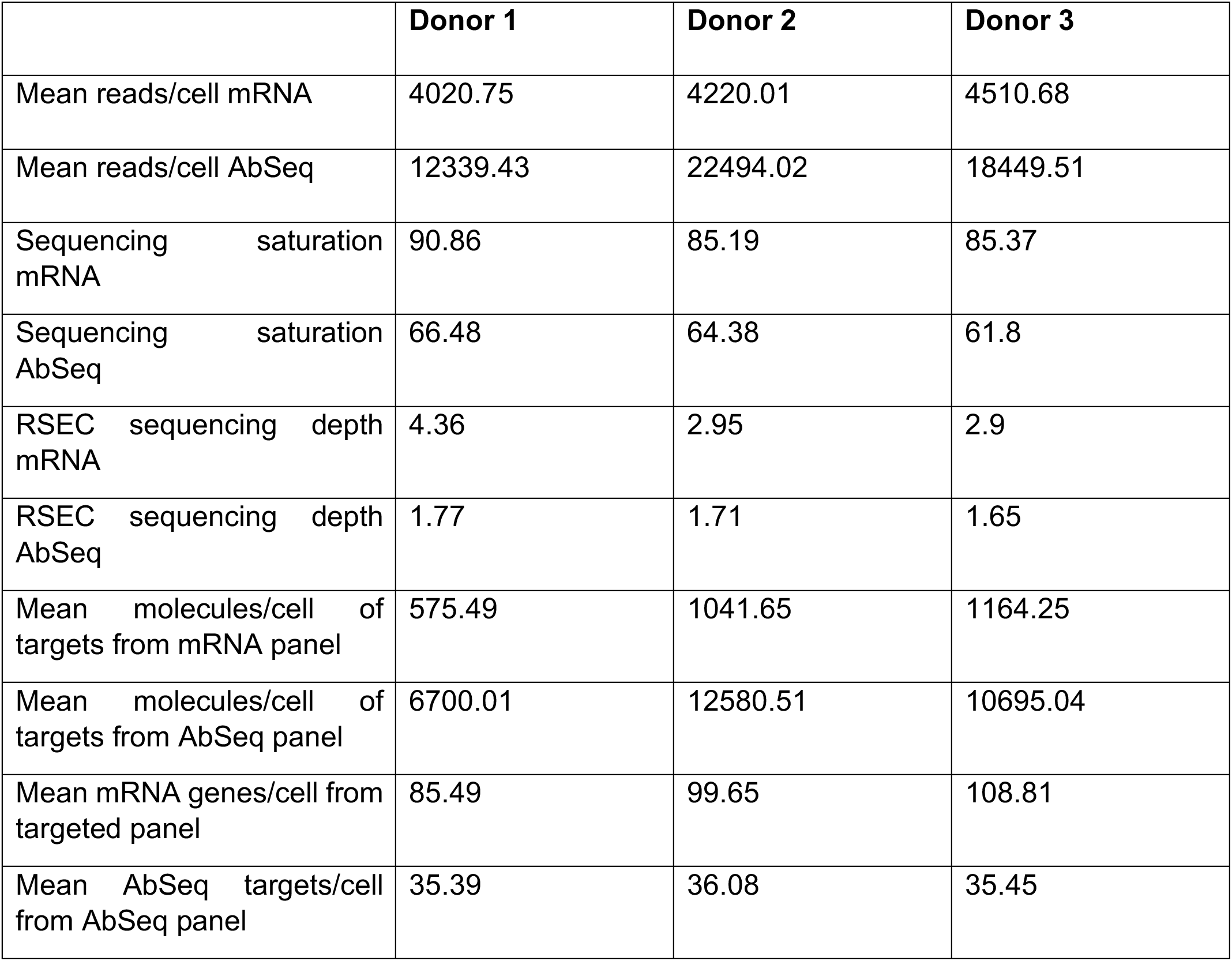
Sequencing metrics.

|  | **Donor 1** | **Donor 2** | **Donor 3** |
| --- | --- | --- | --- |
| Mean reads/cell mRNA | 4020.75 | 4220.01 | 4510.68 |
| Mean reads/cell AbSeq | 12339.43 | 22494.02 | 18449.51 |
| Sequencing saturation mRNA | 90.86 | 85.19 | 85.37 |
| Sequencing saturation AbSeq | 66.48 | 64.38 | 61.8 |
| RSEC sequencing depth mRNA | 4.36 | 2.95 | 2.9 |
| RSEC sequencing depth AbSeq | 1.77 | 1.71 | 1.65 |
| Mean molecules/cell of targets from mRNA panel | 575.49 | 1041.65 | 1164.25 |
| Mean molecules/cell of targets from AbSeq panel | 6700.01 | 12580.51 | 10695.04 |
| Mean mRNA genes/cell from targeted panel | 85.49 | 99.65 | 108.81 |
| Mean AbSeq targets/cell from AbSeq panel | 35.39 | 36.08 | 35.45 |

**Supplemental Figure 1.**
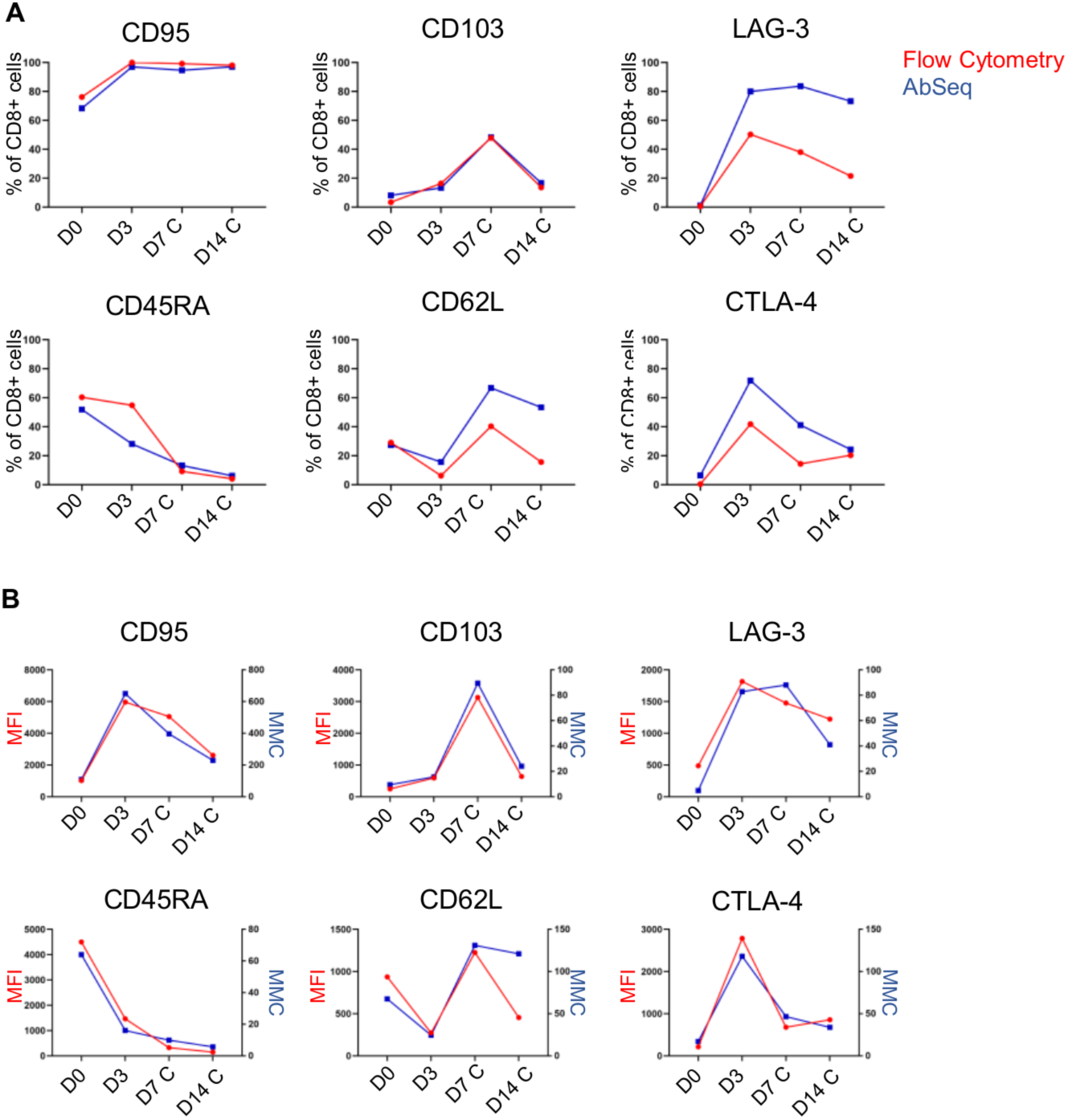
Comparison of protein expression analysis via flow cytometry and AbSeq. (A) Percentage of CD8^+^ cells expressing CD95, CD103, LAG-3, CD45RA, CD62L and CTLA-4 at day 0 (D0), day 3 (D3), day 7 chronic (D7 C) and day 14 chronic (Day14 C) of chronic stimulation using either flow cytometry (red trace) or AbSeq (blue trace). (B) Expression of CD95, CD103, LAG-3, CD45RA, CD62L and CTLA-4 on CD8^+^ T cells at day 0 (D0), day 3 (D3), day 7 (D7 C) and day 14 (Day14 C) of chronic stimulation. Mean fluorescence intensity (red trace, left y axis) and mean molecular count (blue trace, right y axis) were used to measure relative antigen expression levels using flow cytometry and AbSeq, respectively. This side-by-side analysis was performed using aliquots of cryopreserved cells derived from the same donor (Donor 1), and collected at the indicated time points. Flow cytometry data were down-sampled in order to analyze the same number of cells as AbSeq.

**Supplemental Figure 2.**
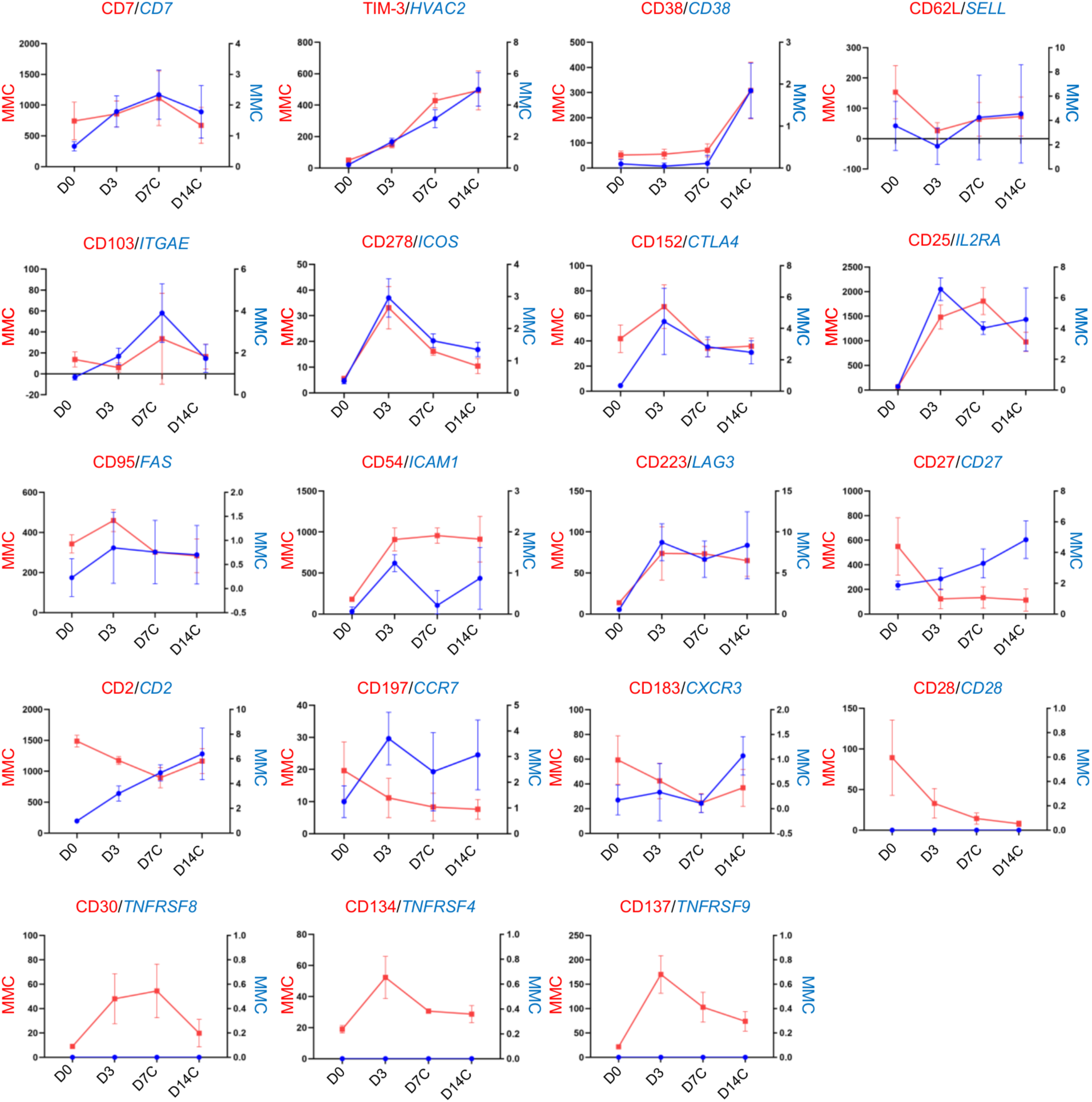
Correlation between gene and protein expression at the single cell level. (A) Kinetic expression of genes and corresponding proteins measured as mean molecular count (MMC; red, left y axis = protein expression; blue, right y axis = gene expression) at day 0 (D0), day 3 (D3), day 7 chronic (D7 C) and day 14 (D14 C) of chronic stimulation. The analysis was performed on three donors. Data are represented as mean ± standard deviation.

